# Motion regions are modulated by scene content

**DOI:** 10.1101/255406

**Authors:** Didem Korkmaz Hacialihafiz, Andreas Bartels

**Author notes:** Corresponding author: Andreas Bartels Vision and Cognition Lab Centre for Integrative Neuroscience, University of Tübingen, Otfried-Müller-Str. 25 72076 Tübingen Germany.

## Abstract

Creating a stable perception of the world during pursuit eye movements is one of the everyday roles of visual system. Some motion regions have been shown to differentiate between motion in the external world from that generated by eye movements. However, in most circumstances, perceptual stability is consistently related to content: the surrounding scene is typically stable. However, no prior study has examined to which extent motion responsive regions are modulated by scene content, and whether there is an interaction between content and motion response. In the present study we used a factorial design that has previously been shown to reveal regional involvement in integrating efference copies of eye-movements with retinal motion to mediate perceptual stability and encode real-world motion. We then added scene content as a third factor, which allowed us to examine to which extent real-motion, retinal motion, and static responses were modulated by meaningful scenes versus their Fourier scrambled counterpart. We found that motion responses in human motion responsive regions V3A, V6, V5+/MT+ and cingulate sulcus visual area (CSv) were all modulated by scene content. Depending on the region, these motion-content interactions differentially depended on whether motion was self-induced or not. V3A was the only motion responsive region that also showed responses to still scenes. Our results suggest that contrary to the two-pathway hypothesis, scene responses are not isolated to ventral regions, but also can be found in dorsal areas.

## Introduction

The visual system encounters different types of motion in dynamic scenes every day and processes visual scenes. Prior studies on motion processing are mostly based on abstract stimuli, like gratings or random dot displays (Born & Bradley, 2005; Boussaoud, Ungerleider, & Desimone, 1990; Erickson & Thier, 1991; Galletti & Fattori, 2003; Goossens, Dukelow, Menon, Vilis, & van den Berg, 2006; Gu, DeAngelis, & Angelaki, 2007; Huk, Dougherty, & Heeger, 2002; Maciokas & Britten, 2010; Smith, Wall, Williams, & Singh, 2006). However, most of these previous studies on motion processing have not used natural scenes, except for one that differentiated self-motion and object motion during movie viewing (Bartels, Zeki, & Logothetis, 2008). Hence, little is known how scene-content influences motion processing and whether these motion regions’ responses are modulated by natural scene content.

Compared to V5+/MT+, which is a well-studied, low-level region in the motion processing hierarchy (Dubner & Zeki, 1971; Zeki et al., 1991), higher-level motion responsive regions such as V6, V3A, or CSv are involved in processing of more complex motion, for instance self-induced visual motion (Fischer, Bulthoff, Logothetis, & Bartels, 2011), integration of self motion cues with vestibular signals (Chowdhury, Takahashi, DeAngelis, & Angelaki, 2009; Gu et al., 2007), or full-field flow compatible with ego-motion (Arnoldussen, Goossens, & van den Berg, 2011; Goossens et al., 2006).

Previous studies showed content related responses in motion processing regions. For instance, human V5/MT has object responses and shows an interaction between object content and motion (Kourtzi, Bulthoff, Erb, & Grodd, 2002; Kourtzi & Kanwisher, 2000). V5/MT, as well as another motion region V3A, were shown to have object selective, size dependent and viewpoint dependent responses (Konen & Kastner, 2008). Further, V3A is shape sensitive (Denys et al., 2004; Grill Spector, Kushnir, Edelman, Itzchak, & Malach, 1998) and involved in form processing (Schira, Fahle, Donner, Kraft, & Brandt, 2004). Importantly, V3A was shown to have a role in scene segmentation (Scholte, Jolij, Fahrenfort, & Lamme, 2008). Additionally, another higher-level motion area V6, that is neighbouring V3A, analyses form and movement in visual field (Galletti et al., 2001). Despite the number of studies pointing out responses related to shape, form or object processing, the effect of scene content on motion regions is not truly known yet.

In this fMRI study, we were interested in whether motion responsive regions modulated by scene content and if they are, how their motion responses depend on scene content. To investigate these questions, we designed stimuli according to a previously established 2 × 2 factorial design with the factors objective motion (on/off) and pursuit (on/off) and this design led us to distinguish objective ‘real’ motion from retinal motion during smooth pursuit eye movements (Fischer, Bulthoff, Logothetis, & Bartels, 2012). In addition, we also added another factor for scene content (gray scale landscape and cityscape scenes or Fourier scrambled versions of these scenes). To balance attention across all conditions, participants performed a central character-matching task at all times. We performed GLM whole-brain analyses as well as region of interest (ROI) analyses. Motion responsive regions were identified using a dedicated localizer scan.

We found that all motion responsive regions tested in this study, namely V5+/MT+, V3A, V6 and CSv, indeed showed scene responses in context of motion. More interestingly, only V3A was responsive to still scenes. The responses in V5+/MT+ were modulated by scene content during both objective and retinal motion, whereas in V3A and V6, the responses were only modulated by scene content during retinal motion. In CSv, although significant responses were present for scenes, we did not observe any interaction between motion and scene content. These results show the importance of naturalistic stimuli use in understanding the visual system and its adaptation to everyday natural scenes.

## Materials and Methods

### Participants

17 healthy participants with normal or corrected-to normal vision (9 female, 1 left-handed, between the age of 20 and 36) gave written informed consent before participating in this study. The study was approved by ethics committee of the University Hospital of Tübingen. All participants were given instructions about the experiment and the task before going into the scanner.

### Experimental Setup

This study consisted of one main experiment, one functional localizer for identifying motion regions V5+/MT+, V3A, V6 and CSv and one structural scan.

The gamma corrected visual stimuli was back-projected onto a screen via a projector outside the scanner room. The visual field of the screen was 19 × 15 visual degrees.

The main experiment was programmed using Psychtoolbox-3 (Brainard 1997, Kleiner, Brainard et al. 2007) whereas the functional localizer experiment that is used for localizing motion regions was prepared using Cogent Graphics v.1.29 developed by John Romaya at the Wellcome Department of Imaging Neuroscience (http://www.vislab.ucl.ac.uk/cogent.php). All stimuli was then presented using a windows PC and MATLAB 7.10.0 (The Mathworks, Natick, MA, 2010) (MATLAB, 2010).

### Main Experiment

The main experiment consisted of eight conditions forming a 2 × 2 × 2 factorial design with the factors objective motion (on/off), pursuit (on/off) and scene (on/off), resulting in the eight conditions (Figure 1). The first two factors were described in a previous study (Fischer et al., 2012).

We picked 32 images of outdoor scenes (both landscape and cityscape) and converted them to grayscale, identical to the images used in one of our previous studies (Korkmaz Hacialihafiz & Bartels, 2015). These grayscale images and their phase-scrambled versions composed the stimuli. In order to balance horizontal inequalities in the images, half of the images were left right flipped duplicates of the other half of the images. All images were adjusted so that they had equal contrast and luminance (luminance: 144 cd/m2, contrast: 32.4 cd/m2 root-mean-square (RMS) contrast, resulting in an average Michelson contrast of 0.90 ± 0.09). We used images that were large enough to give a feeling of moving across the screen.

In order to construct phase-scrambled versions of the images, Fourier transformation was applied and images were reconstructed with random phases. This resulted in preservation of low-level features of the image such as luminance, contrast and spatial frequencies while removing scene content.

The stimuli were presented in a block design. Each run consisted of 33 blocks per run. Each block lasted 12 seconds. The eight conditions were pseudorandomized and back matched so that each condition was preceded by all the other conditions equally. The sequence that allowed this back matching was then divided into two runs. We did this twice in order to obtain 4 runs in total. Each participant took part in 4 runs in total. This way, each condition preceded by each condition in equal frequency across two runs. Moreover, one additional block was added to the beginning of each run in order to initially counterbalance the first block. The images were randomly chosen for each block and only one image was used for an entire block. Stimuli followed a sine trajectory, extending across 4 cycles per block, in order to have a smooth horizontal motion. The velocity varied between 0 and 3.08 deg/s, yielding a mean velocity of 2.53 deg/s and the motion extended up to 1.98 visual degrees in each direction. The starting direction of motion was pseudorandomized and counterbalanced across runs. Each run started with 6.9 seconds of gray screen with fixation and ended with 10 seconds of gray screen with fixation (luminance of gray screen(s): 144 cd/m2). Each run lasted a total of 412.9 seconds. During the experiment, there was a gray fixation disk (width: 0.74 deg, luminance: 282 cd/m2) present at all times on the centre of the screen, with the fixation task described below.

### Fixation Task

There was a 1-back character-matching task at all times in both main experiment and localizer experiment to provide fixation and balanced attention of participants. On the fixation disk, a randomly chosen alphabetical character (a-z) was presented for 1 second each with 83 ms blank intervals in between. Every 3 to 8 presentations, a repetition of the presented character occurred, where participants were required to press a button once they see the repetition. The timings of button presses were recorded and used in GLM analyses as a regressor of no interest.

**Figure 1:**
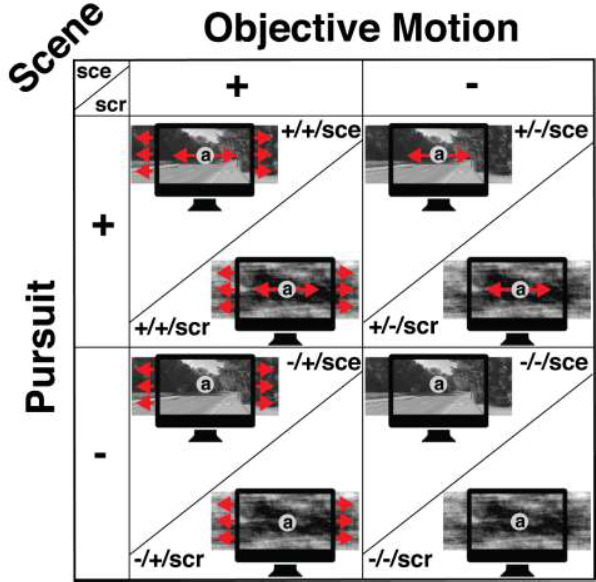
Stimuli. Eight conditions were obtained by a 2 × 2 × 2 factorial design with the following factors: objective motion (on/off), pursuit (on/ off) and scene (scene/ scrambled). . In the “±/±” notation, the first position refers to pursuit, the second to objective motion. “+” refers to presence and “-“ to absence. Objective motion was horizontal motion of the background image (scenes or scrambled images) and pursuit was horizontal motion of the fixation disk. There was a one-back character-matching task inside the fixation disk (shown larger for illustration).

### Functional Localizer

Visual stimuli for the functional motion localizer consisted of random dot patterns. 7 conditions were present in this localizer and each condition was presented 6 times during the session in a pseudorandom history-matched manner. The conditions were as following: 3D full-field motion (coherent motion), random motion, right and left hemifield 3D full-field motion (left or right 1/3rd of the screen), 2D lateral motion with synched pursuit (coherent motion), smooth pursuit with static background and static dots with static fixation task. Each block lasted 12 seconds. Participants performed a 1-back character-matching task, identical to the task in main experiment. Motion areas V3A, V6 and CSv were localized as described previously (Fischer et al., 2011, 2012) and V5+/MT+ was localized using random motion.

Due to technical problems, we could not use right and left hemifield 3D full-field motion conditions to define MT and MST separately, as established previously. Instead, V5+/MT+ was localized using responses to random dot-motion versus static dots. CSv was localized as described previously, using responses to full-field coherent motion with coherently moving fixation dot versus random dot-motion with still fixation(Fischer et al., 2011). V6 was also localized using the same contrast. V3A was localized using responses to coherent 2D motion versus moving fixation dot on static background consisting of dots. All regions are defined using an individual p-value.

### Data Acquisition

T2* weighted functional images were acquired using a 64-channel phased-array head coil in a Siemens Magnetom PRISMA 3T scanner (Siemens, Erlangen, Germany). The voxel size was 3 × 3 × 3 mm3 and TR was 2.3 seconds, while TE was 35 ms and flip angle was 79°. The images included 32 slices, in an ascending order. In order to allow T1 equilibration, the first 3 volumes of data (the first 4 volumes for motion localizer) were discarded. We also collected anatomical images for each participant using T1-weighted images (1 × 1 × 1 mm3 resolution).

### FMRI Data Preprocessing and Statistical Analysis

SPM5 toolbox in MATLAB 7.10.0 was used in order to preprocess functional images with the following steps: reslicing and realignment, followed by coregistration of the structural image to the mean functional image, normalization of the data to the Montreal neurological institute (MNI) space and finally spatial smoothing with 6 mm full-width at half maximum Gaussian kernel for single participants and 12 mm for group level analyses.

Data of each participant were analysed separately using the GLM (general linear model) in SPM5. We modelled each condition and button presses, as well as regressors of no interest, which were six motion realignment parameter series and one additional regressor for global signal variance (Desjardins, Kiehl, & Liddle, 2001; Van Dijk et al., 2010). The global signal variance regressor was orthogonalized to the conditions of interest. The data were high pass filtered using a cut-off value of 128s. In addition, the beta images from the first level GLMs of each participant were used for group level analyses.

The ROI analyses were done by defining ROIs using independent localizer for each participant separately and then extraction of mean beta values for each ROI and for each participant. We used MarsBaR toolbox in order to define ROIs (Brett, Anton, Valabregue, & Poline, 2002). Beta values were range normalized between 0 and 1 for each ROI and participant separately. For 4 runs and 8 conditions, the minimum of all these 32 beta values were subtracted from all 32 beta values and then all of them were divided by maximum of these 32 beta values, for each participant and each ROI separately. Repeated measures ANOVAs, as well as paired t-tests were conducted in order to analyse the effects of conditions using statistical analysis software IBM SPSS Statistics version 22.0.

Mauchly’s sphericity test results were considered for the definition of violation of sphericity and Greenhouse-Geisser correction was used in case of violation of sphericity.

### Eye Tracking

Eye tracking of participants during the main experiment was done using an infrared camera based eye tracker system (Eye-Trac 6; Applied Science Laboratories). The steps of preprocessing included blink removal, smoothing of x and y positions using a running average window of 200 milliseconds. We calculated the fixation accuracy by the root mean square error of actual eye position relative to the fixation disk for each condition across participants and runs. Repeated measures ANOVAs were facilitated in order to analyse eye-tracking data

## Results

After independently localizing motion responsive regions V5+/MT+, V3A, V6 and CSv, we analysed their responses to scene content. We localized V5+/MT+ in 30 hemispheres, V3A in 29 hemispheres, V6 in 28 hemispheres and CSv in 25 hemispheres. For all ROIs, raw and normalized mean beta responses are shown separately in figure 2.

**Figure 2:**
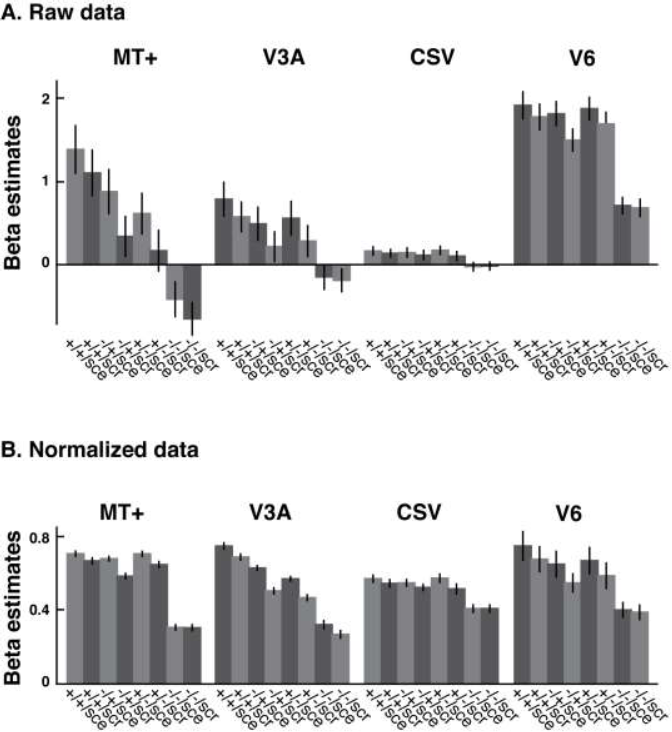
Responses to all conditions across ROIs. (A) Raw beta estimates in motion ROIs V5+/MT+, V3A, CSv and V6. (B) Normalized beta estimates (see methods). In the “±/±” notation, the first position refers to pursuit, the second to objective motion. “+” refers to presence and “-“ to absence. scr: scramble, sce: scene. Plots show mean ± standard error of mean (SEM).

A 4 × 2 × 8 repeated-measures ANOVA with the factors ROI, hemisphere, planar motion, pursuit and scene was conducted in order to test for hemisphere effect. For each ROI we pooled data from both hemispheres since there was no effect of hemisphere (F (1,5) = 0.209, p = 0.667) or any interactions including hemisphere as a factor (hemisphere and ROI: F (6,30) = 0.721, p = 0.636, hemisphere and planar motion: F (1,5) = 0.280, p = 0.620, hemisphere and pursuit: F (1,5) = 0.212, p = 0.664, ROI, hemisphere and planar motion: F (6,30) = 0.643, p = 0.695, ROI, hemisphere, planar motion and pursuit: F (6,30) = 0.980, p = 0.456, hemisphere and scene: F (1,5) = 0.014, p = 0.911, ROI, planar motion, hemisphere and scene: F (6,30) = 0.288, p = 0.938, hemisphere, pursuit and scene: F (1,5) = 1.526, p = 0.272 and hemisphere, ROI, planar motion, pursuit and scene: F (6,30) = 1.172, p = 0.347).

Next, we analysed scene responses in these ROIs and then we analysed content related motion responses, meaning the interactions between scene and different motion types in all ROIs separately.

### Scene responses

We tested scene responses of all ROIs using paired t-tests, using the contrast for all conditions with scenes compared to all conditions with scramble images. All ROIs had a significant scene response (V5+/MT+: t (29) = 8.44, p = 0.11 * 10-7, V3A: t (28) = 7.99, p = 0.32 * 10-7, CSv: t (24) = 2.2, p = 0.038, V6: t (27) = 5.99, p = 0.4 * 10-5, all Bonferroni-Holm corrected for 4 comparisons). Either scenes themselves or motion responses could drive the scene responses in these regions. In order to investigate this, we compared scene versus scramble during still ((-/-/+) versus (-/-/-)) responses using paired t-tests on each motion responsive ROI separately. Only V3A showed a significant difference between responses to scenes and responses to scrambled images during still (t (28) = 4.41, p = 0.0006, Bonferroni-Holm corrected for four comparisons). This difference was not present in other motion responsive regions we investigated (V6 (t (27) = 0.87, p = 0.39), V5+/MT+ (t (29) = 0.69, p = 0.49) and CSv (t (24) = 0.36, p = 0.72). Figure 3A shows scene responses of all ROIs. Next, we tested whether V3A could be differentiated from other regions in its responses to still scenes. Indeed, V3A can be differentiated from all regions by its scene responses (V3A vs. V5+/MT+: t (27) = 2.47, p = 0.02, V3A vs. V6: t (25) = 2.61, p = 0.03, V3A vs. CSv: t (22) = 3.36, p = 0.009, all corrected using Bonferroni-Holm correction for 3 comparisons). Figure 3B shows responses to still scenes in all ROIs.

We also tested scene versus scramble responses during background motion with eye fixation ((-/+/sce) vs. (-/+/scr)). As seen in figure 3C, V5+/ MT+ (t (29) = 8.11, p = 0.25 * 10-7), V3A (t (28) = 6.72, p = 0.81 * 10-6), and V6 (t (27) = 4.27, p = 0.0004) significantly responded to scenes with background motion during fixation, whereas CSv did not (t (24) = 1.08, p = 0.29) (Corrected using Bonferroni-Holm correction for four comparisons).

**Figure 3:**
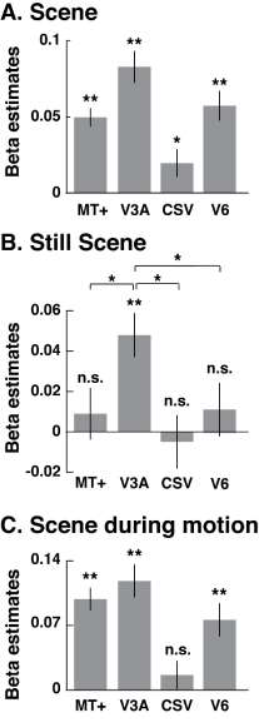
Scene responses in V5+/MT+, V3A, V6 and CSv. (A) Main effect of scenes. (B) Responses to still scenes vs. still scramble images. (C) Responses to moving scenes vs moving scramble images during fixation (-/+/sce vs. -/+/scr). **: p < 0.001, *: p < 0.05. Bonferroni-Holm corrected. Plots show mean ± standard error of mean (SEM).

### Content effect on motion responses

Next, we tested the interactions between motion and scene content in the regions of interests.

First, we tested objective motion responses during scenes, during scrambled images and their interaction. As expected, in all regions there were significant objective motion responses during scenes (V5+/MT+: t (29) = 15.51, p = 0.17 * 10-13, V3A: t (28) = 11.95, p = 0.18 * 10-10, V6: t (27) = 7.59, p = 0.26 * 10-6, CSv: t (24) = 5.2, p = 0.13 * 10-3) and scrambled images (V5+/MT+: t (29) = 10.33, p = 0.29 * 10-9, V3A: t (28) = 11.78, p = 0.23 * 10-10, V6: t (27) = 9.7, p = 0.22 * 10-8, CSv: t (24) = 5.74, p = 0.42 * 10-4). However, there was a significant interaction between objective motion and scene content only in V5+/MT+ (t (29) = 2.94, p = 0.024), although in V3A and V6, there was a trend for higher objective motion responses during scenes compared to during scramble (All corrected for 12 comparisons using Bonferroni-Holm correction).

Next, we tested retinal motion responses during scenes, scrambles and their interaction. All regions showed significant retinal motion responses during scenes (V5+/MT+: t (29) = 15.28, p = 0.25 * 10-13, V3A: t (28) = 5.24, p = 0.14 * 10-3, V6: t (27) = 4.58, p = 0.65 * 10-3, CSv: t (24) = 4.94, p = 0.43 * 10-3). Interestingly, only V5+/MT+ (t (29) = 9.99, p = 0.75 * 10^-9^) and CSv (t (24) = 3.63, p = 0.005) had significant retinal motion responses during scramble images while V3A (t (28) = 0.49, p = 0.627) and V6 (t (27) = 1.28, p = 0.211) did not. All regions except CSv showed a significant interaction between retinal motion and scene content (V5+/MT+: t (29) = 4.74, p = 0.41 * 10-3, V3A: t (28) = 4.03, p = 0.0024, V6: t (27) = 3.06, p = 0.02, CSv: t (24) = 1.57, p = 0.129) (All corrected for 12 comparisons using Bonferroni-Holm correction).

**Figure 4.**
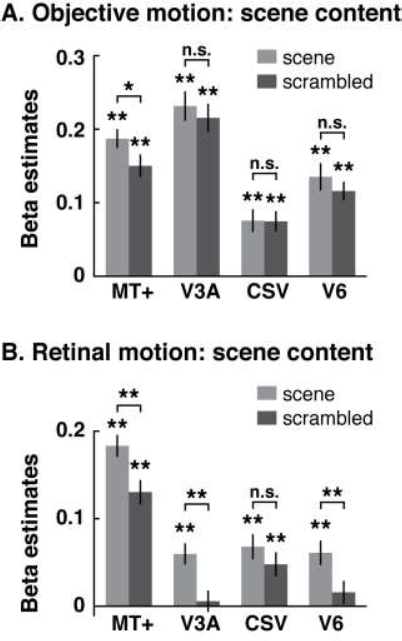
Objective and retinal motion preferences across ROIs. (A) Objective motion responses, shown for scenes and scrambled backgrounds separately and their interactions. (B) Retinal motion responses, shown for scenes and scrambled backgrounds separately and their interactions. **: p < 0.001, *: p < 0.05, Bonferroni-Holm corrected. Plots show mean ± standard error of mean (SEM).

### Whole brain Analyses

Since V3A is responsive to still scenes, we wanted to check if it overlaps with scene responsive areas in the group level analysis. We first calculated the contrast still scenes vs. still scramble images in group level, with p < 0.05 uncorrected. Next, we defined V3A using the previously established contrast (Fischer et al., 2012); objective vs. retinal motion during both scenes and scramble images in the group level using p < 0.001 uncorrected. Figure 5 shows the overlap between these two contrasts.

**Figure 5:**
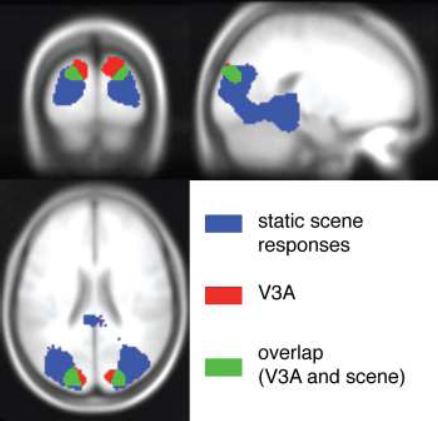
Whole brain results showing overlap between V3A and scene responsive areas. The contrasts were created using (a) still scene vs still scrambled images, shown by blue and (b) objective versus retinal motion to define V3A (during both scenes and scramble images), shown by red and their interaction is shown by green.

### Behaviour Data

Participants performed a character back-matching task during the main experiment. The mean correct response rate was 0.85 ± 0.07 (mean ± std), whereas mean response time was 0.57 ± 0.14 s (mean ± std). We analysed response time data using a 3-way repeated measures ANOVA with the factors objective motion, pursuit and scene. There were only significant effects of pursuit (F (1,16) = 5.06, p = 0.039) but no other main effects (objective motion: F (1,16) = 0.005, p = 0.95; scene: F (1,16) = 0.39, p = 0.54), nor interactions (objective and pursuit (i.e., retinal motion): F (1,16) = 0.93, p = 0.35; objective and scene: F (1,16) = 0.00029, p = 0.99; pursuit and scene F (1,16) = 1.58, p = 0.23; objective, pursuit and scene: F (1,16) = 0.98, p = 0.34).

### Eye tracking Data

Eye tracking data were collected for each participant during scanning and were preprocessed as described in methods. Following preprocessing, RMSE of eye position relative to the fixation disk was calculated and these RMSE were used for calculation and comparison of fixation accuracy. The average RMSE across participants and runs and conditions is 1.54 ± 0.73 deg (mean ± std). We used 2 × 2 × 2 repeated measures ANOVA with factors objective motion, pursuit and scene to analyse the RMSE data. There was an effect of pursuit (F (1, 67) = 682.38, p = 0.75 * 10-36), but there were no effects of objective motion (F (1, 67) = 0.06, p = 0.82) or scene (F (1, 67) = 0.0001, p = 0.99). Moreover, there were no interaction between objective motion and pursuit (which gives us retinal motion) (F (1, 67) = 0.28, p = 0.56), between objective motion and scene (F (1, 67) = 0.14, p = 0.71), between pursuit and scene (F (1, 67) = 2.8, p = 0.099) and between objective motion, pursuit and scene (F (1, 67) = 0.16, p = 0.69).

## Discussion

Here, we sought answers to the following questions: are motion responsive regions modulated by scene content and if they are, how do their responses to objective and retinal motion change with scene content modulation? We used a 2 × 2 × 2 factorial design with the factors being real world scenes versus their scrambled versions, horizontal panning motion and smooth pursuit eye movements. We chose horizontal panning motion and pursuit eye movements, as they are frequently found and are natural in daily life. We examined well-known motion responsive regions V5+/MT+, V3A, V6, and CSv.

We found that the motion responses of all motion regions showed a scene preference, whereas only V3A also responded significantly to still scene images compared to still scrambles. Moreover, V5+/MT+, V3A and V6 showed motion and scene content interaction whereas this was absent in the responses of CSv.

### Scene responses in V3A

V3A was the only region with significant scene responses even in the absence of any type of motion. V3A is an intermediate-tier, retinotopic region with relatively large receptive field size and representing both upper and lower visual fields (Tootell et al., 1997). So, it is unlikely that responses to static scenes could be driven by possible local visual field irregularities between scenes and scrambled images. Human V3A is neighbouring a scene responsive region, occipital place area (OPA), which is also known as transverse occipital sulcus (TOS) (Nasr et al., 2011). Interestingly, some recent studies have shown that OPA partially overlaps with V3A (Bettencourt & Xu, 2013; Silson, Groen, Kravitz, & Baker, 2016). Additionally, a recent study found that V3A is connected to ventral regions (hV4/ VO-1) via a major white matter pathway in human brain, namely vertical occipital fasciculus (VOF) (Takemura et al., 2015). More research is needed to understand the exact role of V3A in scene processing.

Although V3A is known as a motion-processing region, its exact role is still under investigation. Previous studies have shown that V3A processes three-dimensional structure and depth cues as well as shapes defined by colour or motion (Georgieva, Peeters, Kolster, Todd, & Orban, 2009; Paradis et al., 2000; Self & Zeki, 2005). Other findings include that V3A has a role in shape integration, contributes to contour integration, is modulated by context (Aspell, Wattam-Bell, Atkinson, & Braddick, 2010; Schira et al., 2004), has shape sensitive responses and is involved in object processing (Denys et al., 2004; Grill Spector et al., 1998) and has a role in not only in boundary detection, but also in scene segmentation and texture segregation (together with other early level visual areas) (Kastner, De Weerd, & Ungerleider, 2000; Scholte et al., 2008). The responses to still scenes in V3A shown here could be driven by continuous contours, which exist in scene images but absent scrambled images, or by higher-level image features such as shapes or objects within the scenes.

Our results also showed an interaction between scene and retinal motion in V3A, meaning that V3A’s scene responses were higher during motion. Interestingly, while V3A showed significant responses to retinal motion during scenes, it did not show any significant retinal motion responses during scrambled images. We believe this could be driven by viewpoint changes across scenes, since in a previous study, V3A, as well as MT, was shown to have viewpoint specific object selective responses (Konen & Kastner, 2008).

V3A has many connections with parietal regions and is thought to provide information about object location and motion to these regions. This information is probably used by parietal regions during self-object interaction such as reaching and grasping. While encoding object location and motion, V3A is involved in depth perception and having a representation of visual scenes could be useful with this role of V3A.

### Content-dependent responses in motion processing regions

V5+/MT+ showed significant interaction between scene and objective motion and between scenes and retinal motion, meaning that, it responded more to motion (both objective and retinal motion) when there were scenes in the background, compared to when there were scrambled images in the background. It is unlikely that these responses are due to differences of spatial frequency in lower and upper parts of the visual scene images since human V5/MT and MST show no upper-lower visual field bias (Kolster, Peeters, & Orban, 2010). In addition to its well-known motion related responses such as responses to optic flow, speed and direction selective responses, V5/MT also shows object and shape sensitivity as well as viewpoint specific object responses (Denys et al., 2004; Kolster et al., 2010; Konen & Kastner, 2008; Kourtzi et al., 2002; Kourtzi & Kanwisher, 2000). Our results are parallel to the aforementioned findings regarding content effect in V5+/MT+.

Interestingly, V6 showed scene related responses during retinal-motion. Previous studies on V6 showed its involvement in optic flow responses and processing of self-motion related cues (Cardin & Smith, 2010; Fischer et al., 2012; Pitzalis et al., 2010). Additionally, V6 contains representations of both upper and lower visual field (Pitzalis et al., 2006). V6 is highly connected to parietal regions and its visuomotor neighbour V6A. V6A and parietal regions are particularly interested in reaching and grasping. In relation to this, a study showed that V6 is more responsive to near visual field compared to far visual field, and suggested that these responses are related to object locations for reaching (Quinlan & Culham, 2007). It is possible that scene content responses are also related to distance encoding in V6. Thus, motion during scene generates more response compared to motion during scrambled images.

CSv only showed scene responses but no interaction or no scene response during motion or no motion. It is possible that scene responses in CSv are modulated by eye movements, but since pursuit condition is not well controlled, we did not investigate pursuit related responses in this study. CSv is located in posterior cingulate cortex and has been shown to contain information about heading direction during self-motion (Furlan, Wann, & Smith, 2014). CSv also showed vestibular responses (Smith, Wall, & Thilo, 2012). Furthermore, CSv has been shown to integrate eye movements with retinal motion (Fischer et al., 2011). Cingulate sulcus has been previously shown to have place category responses (Epstein & Higgins, 2007). Natural scene images, as we used in this experiment, resembles everyday experience for the visual cortex in a way that it provides realistic input (compared to scrambled images) and perceptually, provides more important cues regarding heading and self-motion. So, in this context, CSv responses could be explained by more engagement of CSv during eye movements in natural scenes since this resembles perceptually more natural input regarding self-motion extraction or heading direction.

Our result showing scene related higher activation in every motion responsive region is rather interesting. Scene related responses can be used during depth perception via textures or perspective whereas this is absent in scrambled images. Previous studies showed that natural stimuli are preferred by the visual cortex (Kayser, Kording, & Konig, 2004). However, more studies on contextual effects of natural scene stimuli on motion processing regions would provide a better insight about the effects seen here.

### Low-level versus high-level interpretations

One can think that the results shown here are due to confound regarding low level differences across scenes and their scrambled versions. Phase scrambling was used in order to conserve low-level image features such as spatial frequency while eliminating contextual effect, thus making the high level aspect of the image (such as scenes as in here, but also used for objects and faces) unrecognizable. Traditionally, applying phase scrambling while keeping luminance and contrast equal across images and their scrambled versions is thought to result in no response from early visual cortex but engage higher level regions’ (or ventral regions’) responsiveness. However, a number of studies raised concerns about using phase-scrambled images in this way. For instance, a comparison of the contrasts of natural scenes and their scrambled counterparts resulted in higher number of peaks in the histogram of natural images (Dumoulin, Dakin, & Hess, 2008). Also, scrambling scatters local constructions within the image all over (Kay, Winawer, Rokem, Mezer, & Wandell, 2013). Related to these concerns, modifications to phase scrambling and even different scrambling methods have been proposed (Ales, Farzin, Rossion, & Norcia, 2012; Stojanoski & Cusack, 2014). Hence, it cannot be ruled out that our findings regarding higher scene responses in motion responsive regions might be related to higher-order or localized differences in low-level image features such as lines or contours. Numerous studies have investigated how different level features are processed in human brain. More related to our findings, extra-striate visual cortex has been shown to have contour-based responses to scene images (Dumoulin et al., 2008). Another study showed that a comparison of lines and edges to phase scrambled images created higher responses for lines and edges in most of the visual areas, even early visual cortex (Perna, Tosetti, Montanaro, & Morrone, 2008). For ventral scene-responsive regions, a series of recent studies provided evidence that low-level features may account for their previously reported preference to certain high-level categories (Nasr, Echavarria, & Tootell, 2014; Nasr & Tootell, 2012; Rajimehr, Devaney, Bilenko, Young, & Tootell, 2011). These studies argued that low-level features typically associated to scenes, such as cardinal orientations, rectilinearity, and high spatial frequencies alone selectively activate PPA. However, more recent evidence showed that even when all low-level features are controlled for, PPA still prefers high-level interpretations of features perceived as spatial arrangements (Bryan, Julian, & Epstein, 2016; Schindler & Bartels, 2016). Clearly, further more detailed studies are needed in order to clarify the underlying mechanism of scene related responses in motion responsive regions.

## Conclusion

In conclusion, V5+/MT+, V3A, V6 and CSv had content effect due to scene content during motion. V3A also had scene responses during still scenes. These results contribute to our understanding of how V5+/MT+, V3A, V6 and CSv responses are modulated by scene content.

These results support the view that unlike the traditional theories about completely segregated dorsal ‘what’ and ventral ‘where’ streams, these two pathways functionally interact. Consistent with this view, our results could be interpreted as V3A taking part in analysing the 3D overlay of the visual scenes, which can be useful when calculating the motion of objects in depth. Further studies are needed to investigate the role of motion regions, especially V3A, in detail during scene viewing.

## Acknowledgements

This work was funded by the Centre for Integrative Neuroscience Tübingen through the German Excellence Initiative (EXC307) and by the Max Planck Society, Germany.

## References

Ales, J. M., Farzin, F., Rossion, B., & Norcia, A. M. (2012). An objective method for measuring face detection thresholds using the sweep steady-state visual evoked response. Journal of Vision, 12(10), 18–18. doi: 10.1167/12.10.18

Arnoldussen, D. M., Goossens, J., & van den Berg, A. V. (2011). Adjacent visual representations of self-motion in different reference frames. Proceedings of the National Academy of Sciences of the United States of America, 108, 11668–11673. doi: 10.1073/pnas.1102984108

Aspell, J. E., Wattam-Bell, J., Atkinson, J., & Braddick, O. J. (2010). Differential human brain activation by vertical and horizontal global visual textures. Experimental brain research, 202(3), 669–679.

Bartels, A., Zeki, S., & Logothetis, N. K. (2008). Natural Vision Reveals Regional Specialization to Local Motion and to Contrast-Invariant, Global Flow in the Human Brain. Cerebral Cortex, 18(3), 705–717. doi: 10.1093/cercor/bhm107

Bettencourt, K. C., & Xu, Y. (2013). The Role of Transverse Occipital Sulcus in Scene Perception and Its Relationship to Object Individuation in Inferior Intraparietal Sulcus. Journal of Cognitive Neuroscience, 25(10), 1711–1722. doi: http://dx.doi.org/10.1162/jocn_a_00422

Born, R. T., & Bradley, D. C. (2005). Structure and function of visual area MT Annual Review of Neuroscience (Vol. 28, pp. 157–189). Palo Alto: Annual Reviews.

Boussaoud, D., Ungerleider, L. G., & Desimone, R. (1990). Pathways for Motion Analysis - Cortical Connections of the Medial Superior Temporal and Fundus of the Superior Temporal Visual Areas in the Macaque. Journal of Comparative Neurology, 296(3), 462–495.

Brett, M., Anton, J.-L., Valabregue, R., & Poline, J.-B. (2002). Region of interest analysis using the MarsBar toolbox for SPM 99. Neuroimage, 16(2), S497.

Bryan, P. B., Julian, J. B., & Epstein, R. (2016). Rectilinear edge selectivity is insufficient to explain the category selectivity of the parahippocampal place area. Frontiers in human neuroscience, 10.

Cardin, V., & Smith, A. T. (2010). Sensitivity of human visual and vestibular cortical regions to egomotion-compatible visual stimulation. Cerebral cortex, 20(8), 1964–1973. doi: 10.1093/cercor/bhp268

Chowdhury, S. A., Takahashi, K., DeAngelis, G. C., & Angelaki, D. E. (2009). Does the middle temporal area carry vestibular signals related to self-motion? The Journal of neuroscience: the official journal of the Society for Neuroscience, 29(38), 12020–12030. doi: 10.1523/JNEUROSCI.0004-09.2009

Denys, K., Vanduffel, W., Fize, D., Nelissen, K., Peuskens, H., Van Essen, D., & Orban, G. A. (2004). The processing of visual shape in the cerebral cortex of human and nonhuman primates: a functional magnetic resonance imaging study. J Neurosci, 24(10), 2551–2565.

Desjardins, A. E., Kiehl, K. A., & Liddle, P. F. (2001). Removal of confounding effects of global signal in functional MRI analyses. Neuroimage, 13(4), 751–758. doi: http://dx.doi.org/10.1006/nimg.2000.0719

Dubner, R., & Zeki, S. (1971). Response properties and receptive fields of cells in an anatomically defined region of the superior temporal sulcus in the monkey. Brain Research, 35, 528–532.

Dumoulin, S. O., Dakin, S. C., & Hess, R. F. (2008). Sparsely distributed contours dominate extra-striate responses to complex scenes. Neuroimage, 42(2), 890–901. doi: 10.1016/j.neuroimage.2008.04.266

Epstein, R., & Higgins, J. S. (2007). Differential parahippocampal and retrosplenial involvement in three types of visual scene recognition. Cereb Cortex, 17(7), 1680–1693. doi: http://dx.doi.org/10.1093/cercor/bhl079

Erickson, R. G., & Thier, P. (1991). A Neuronal Correlate of Spatial Stability during Periods of Self-Induced Visual-Motion. Experimental Brain Research, 86(3), 608–616.

Fischer, E., Bulthoff, H. H., Logothetis, N. K., & Bartels, A. (2011). Visual Motion Responses in the Posterior Cingulate Sulcus: A Comparison to V5/MT and MST. Cerebral cortex. doi: 10.1093/cercor/bhr154

Fischer, E., Bulthoff, H. H., Logothetis, N. K., & Bartels, A. (2012). Human areas V3A and V6 compensate for self-induced planar visual motion. Neuron, 73(6), 1228–1240. doi: 10.1016/j.neuron.2012.01.022

Furlan, M., Wann, J. P., & Smith, A. T. (2014). A Representation of Changing Heading Direction in Human Cortical Areas pVIP and CSv. Cerebral Cortex, 24(11), 2848–2858. doi: 10.1093/cercor/bht132

Galletti, C., & Fattori, P. (2003). Neuronal mechanisms for detection of motion in the field of view. Neuropsychologia, 41(13), 1717–1727.

Galletti, C., Gamberini, M., Kutz, D. F., Fattori, P., Luppino, G., & Matelli, M. (2001). The cortical connections of area V6: an occipito-parietal network processing visual information. Eur Neurosci, 13(8), 1572–1588.

Georgieva, S., Peeters, R., Kolster, H., Todd, J. T., & Orban, G. A. (2009). The processing of three-dimensional shape from disparity in the human brain. The Journal of neuroscience: the official journal of the Society for Neuroscience, 29, 727–742. doi: 10.1523/JNEUROSCI.4753-08.2009

Goossens, J., Dukelow, S. P., Menon, R. S., Vilis, T., & van den Berg, A. V. (2006). Representation of Head-Centric Flow in the Human Motion Complex. The Journal of Neuroscience, 26(21), 5616–5627. doi: 10.1523/jneurosci.0730-06.2006

Grill Spector, K., Kushnir, T., Edelman, S., Itzchak, Y., & Malach, R. (1998). Cue-invariant activation in object-related areas of the human occipital lobe. NEURON. Neuron., 21(1), 191–202.

Gu, Y., DeAngelis, G. C., & Angelaki, D. E. (2007). A functional link between area MSTd and heading perception based on vestibular signals. Nature Neuroscience, 10(8), 1038–1047.

Huk, A. C., Dougherty, R. F., & Heeger, D. J. (2002). Retinotopy and Functional Subdivision of Human Areas MT and MST. The Journal of Neuroscience, 22(16), 7195–7205.

Kastner, S., De Weerd, P., & Ungerleider, L. G. (2000). Texture Segregation in the Human Visual Cortex: A Functional MRI Study. Journal of Neurophysiology, 83(4), 2453–2457.

Kay, K. N., Winawer, J., Rokem, A., Mezer, A., & Wandell, B. A. (2013). A Two-Stage Cascade Model of BOLD Responses in Human Visual Cortex. PLoS Comput Biol, 9(5), e1003079. doi: 10.1371/journal.pcbi.1003079

Kayser, C., Kording, K. P., & Konig, P. (2004). Processing of complex stimuli and natural scenes in the visual cortex. Curr Opin Neurobiol, 14(4), 468–473.

Kolster, H., Peeters, R., & Orban, G. A. (2010). The retinotopic organization of the human middle temporal area MT/V5 and its cortical neighbors. The Journal of neuroscience: the official journal of the Society for Neuroscience, 30, 9801–9820. doi: 10.1523/JNEUROSCI.2069-10.2010

Konen, C. S., & Kastner, S. (2008). Two hierarchically organized neural systems for object information in human visual cortex. Nature neuroscience, 11, 224–231. doi: 10.1038/nn2036

Korkmaz Hacialihafiz, D., & Bartels, A. (2015). Motion responses in scene-selective regions. NeuroImage, 118, 438–444. doi: http://dx.doi.org/10.1016/j.neuroimage.2015.06.031

Kourtzi, Z., Bulthoff, H. H., Erb, M., & Grodd, W. (2002). Object-selective responses in the human motion area MT/MST. Nat Neurosci, 5(1), 17–18. doi: http://www.nature.com/neuro/journal/v5/n1/suppinfo/nn780_S1.html

Kourtzi, Z., & Kanwisher, N. (2000). Activation in human MT/MST by static images with implied motion. Journal of Cognitive Neuroscience, 12(1), 48–55.

Maciokas, J. B., & Britten, K. H. (2010). Extrastriate Area MST and Parietal Area VIP Similarly Represent Forward Headings. Journal of Neurophysiology, 104(1), 239–247. doi: 10.1152/jn.01083.2009

MATLAB. (2010). version 7.10.0 Natick, Massachusetts: The MathWorks Inc.

Nasr, S., Echavarria, C. E., & Tootell, R. B. H. (2014). Thinking Outside the Box: Rectilinear Shapes Selectively Activate Scene-Selective Cortex. The Journal of Neuroscience, 34(20), 6721–6735. doi: 10.1523/JNEUROSCI.4802-13.2014

Nasr, S., Liu, N., Devaney, K. J., Yue, X., Rajimehr, R., Ungerleider, L. G., & Tootell, R. B. H. (2011). Scene-selective cortical regions in human and nonhuman primates. The Journal of neuroscience: the official journal of the Society for Neuroscience, 31, 13771–13785. doi: 10.1523/JNEUROSCI.2792-11.2011

Nasr, S., & Tootell, R. B. H. (2012). A cardinal orientation bias in scene-selective visual cortex. The Journal of neuroscience: the official journal of the Society for Neuroscience, 32, 14921–14926. doi: http://dx.doi.org/10.1523/JNEUROSCI.2036-12.2012

Paradis, A. L., Cornilleau-Pérès, V., Droulez, J., Van De Moortele, P. F., Lobel, E., Berthoz, A., … Poline, J. B. (2000). Visual perception of motion and 3-D structure from motion: an fMRI study. Cerebral cortex (New York, N.Y.: 1991), 10, 772–783.

Perna, A., Tosetti, M., Montanaro, D., & Morrone, M. C. (2008). BOLD response to spatial phase congruency in human brain. Journal of Vision, 8(10), 15–15. doi: 10.1167/8.10.15

Pitzalis, S., Galletti, C., Huang, R. S., Patria, F., Committeri, G., Galati, G., … Sereno, M. I. (2006). Wide-field retinotopy defines human cortical visual area v6. J Neurosci, 26(30), 7962–7973.

Pitzalis, S., Sereno, M. I., Committeri, G., Fattori, P., Galati, G., Patria, F., & Galletti, C. (2010). Human v6: the medial motion area. Cerebral cortex, 20(2), 411–424. doi: 10.1093/cercor/bhp112

Quinlan, D. J., & Culham, J. C. (2007). fMRI reveals a preference for near viewing in the human parieto-occipital cortex. NeuroImage, 36(1), 167–187. doi: 10.1016/j.neuroimage.2007.02.029

Rajimehr, R., Devaney, K. J., Bilenko, N. Y., Young, J. C., & Tootell, R. B. H. (2011). The “parahippocampal place area” responds preferentially to high spatial frequencies in humans and monkeys. PLoS biology, 9, e1000608. doi: 10.1371/journal.pbio.1000608

Schindler, A., & Bartels, A. (2016). Visual high-level regions respond to high-level stimulus content in the absence of low-level confounds. NeuroImage, 132, 520–525.

Schira, M. M., Fahle, M., Donner, T. H., Kraft, A., & Brandt, S. A. (2004). Differential Contribution of Early Visual Areas to the Perceptual Process of Contour Processing. Journal of Neurophysiology, 91(4), 1716–1721.

Scholte, H. S., Jolij, J., Fahrenfort, J. J., & Lamme, V. A. F. (2008). Feedforward and recurrent processing in scene segmentation: electroencephalography and functional magnetic resonance imaging. Journal of cognitive neuroscience, 20, 2097–2109. doi: 10.1162/jocn.2008.20142

Self, M. W., & Zeki, S. (2005). The integration of colour and motion by the human visual brain. Cerebral cortex, 15, 1270–1279. doi: http://dx.doi.org/10.1093/cercor/bhi010

Silson, E. H., Groen, I. A., Kravitz, D. J., & Baker, C. I. (2016). Evaluating the correspondence between face-, scene-, and object-selectivity and retinotopic organization within lateral occipitotemporal cortex. Journal of vision, 16(6), 14–14.

Smith, A. T., Wall, M. B., & Thilo, K. V. (2012). Vestibular inputs to human motion-sensitive visual cortex. Cerebral cortex (New York, N.Y.: 1991), 22, 1068–1077. doi: 10.1093/cercor/bhr179

Smith, A. T., Wall, M. B., Williams, A. L., & Singh, K. D. (2006). Sensitivity to optic flow in human cortical areas MT and MST. European Journal of Neuroscience, 23(2), 561–569.

Stojanoski, B., & Cusack, R. (2014). Time to wave good-bye to phase scrambling: Creating controlled scrambled images using diffeomorphic transformations. Journal of Vision, 14(12), 6–6. doi: 10.1167/14.12.6

Takemura, H., Rokem, A., Winawer, J., Yeatman, J. D., Wandell, B. A., & Pestilli, F. (2015). A Major Human White Matter Pathway Between Dorsal and Ventral Visual Cortex. Cerebral Cortex. doi: 10.1093/cercor/bhv064

Tootell, R. B. H., Mendola, J. D., Hadjikhani, N. K., Ledden, P. J., Liu, A. K., Reppas, J. B., … Dale, A. M. (1997). Functional analysis of V3A and related areas in human visual cortex. Neurosci, 17(18), 7060–7078.

Van Dijk, K. R., Hedden, T., Venkataraman, A., Evans, K. C., Lazar, S. W., & Buckner, R. L. (2010). Intrinsic functional connectivity as a tool for human connectomics: theory, properties, and optimization. J Neurophysiol, 103(1), 297–321. doi: http://dx.doi.org/10.1152/jn.00783.2009

Zeki, S., Watson, J. D., Lueck, C. J., Friston, K. J., Kennard, C., & Frackowiak, R. S. (1991). A direct demonstration of functional specialization in human visual cortex. The Journal of Neuroscience, 11(3), 641–649.

